# Strategic Inattention in the Sir Philip Sidney Game

**DOI:** 10.1101/559955

**Authors:** Mark Whitmeyer

## Abstract

Infamously, the presence of honest communication in a signaling environment may be difficult to reconcile with small (relative) signaling costs or a low degree of common interest between sender (beneficiary) and receiver (donor). This paper posits that one mechanism through which such communication can arise is through inattention on the part of the receiver, which allows for honest communication in settings where–should the receiver be fully attentive–honest communication would be impossible. We explore this idea through the Sir Philip Sidney game in detail and show that some degree of inattention is always weakly better for the receiver and may be strictly better. We compare limited attention to Lachmann and Bergstrom’s (1998) notion of a signaling medium and show that the receiver-optimal degree of inattention is equivalent to the receiver-optimal choice of medium.

## 1 Introduction

> I have only one eye–I have a right to be blind sometimes.
>
> …I really do not see the signal.
>
> Admiral Horatio Lord Nelson

One standard way of formally studying information transmission between organisms is the signaling game, in which an agent (the sender) signals to a second player (the receiver), who subsequently takes an action. The literature on such games began with the seminal work of Lewis (1969), who discusses costless (“cheap talk”) communication between parties with completely aligned interests. In his formulation, without frictions or signaling costs, there exist equilibria in which communication occurs. However, in many instances the aims and objectives of the communicating parties are *not* fully aligned. Can communication occur then?

Remarkably, as discovered by Crawford and Sobel (1982), even when the interests of the communicating parties diverge and talk is cheap, some information transmission can occur, provided the interests of the communicating parties do not differ by too much. Unsurprisingly, though, as noted by Spence (1973), heterogeneous signaling costs for the different types of sender can greatly enhance or facilitate communication. This idea is related to the handicap principle in biology (Zahavi (1975)), which states that order to facilitate meaningful communication in situations in which there are conflicts of interest, a cost is necessary. A number of recent works^1^ have subsequently argued that it is not the costly aspect of the signal that is important, *per se*, but rather the signal’s *relative* cost for the bad (dishonest) types. However, the basic point remains: for communication to occur, it must be too costly for bad types to mimic good types.

One thorn in the paw of this theory is the empirical finding that the observed signaling costs in nature may not be sufficiently high to ensure honesty, at least in some paradigmatic settings. Indeed, in recent work, Zollman et al. (2013) state that, “researchers have not always been able to find substantial signal costs associated with putative costly signal systems–despite evidence that these systems do convey, at least, some information among individuals with conflicting interests,” and ask, “What then, are we to make of empirical situations in which signals appear to be informative even without the high costs required by costly signaling models?” Zollman et al. mention that one possible solution to this issue is suggested by a number of works–see, e.g., the papers listed in Footnote 1 above–who illustrate that this issue may be ameliorated by recognizing that costly signals need not be sent on the equilibrium path. That is, it is the high cost of sending a (deviating) signal that keeps the sender types honest.

In contrast, this paper explores an alternative mechanism through which communication can be sustained despite low (relative) signaling costs: *inattention* on the part of the receiver. The setting for this analysis is the well-known signaling game, the discrete Sir Philip Sidney Game (Smith (1991)). The classic formulation of the game is quite straightforward: two players interact, a beneficiary and a donor. The type of the beneficiary, or **sender**, is uncertain and private information for the sender: he is either healthy (with probability 1 − *µ*) or needy (with probability *µ*). The sender is the first mover, and may choose to either cry out and incur a cost of *c* > 0, or stay silent and incur no cost. Following this action (henceforth referred to as a *signal*), the donor, or **receiver**, observes the signal before choosing whether to donate a resource and incur a cost of *d* > 0 or do nothing and incur no cost.

Should a sender receive a donation, his probability of survival is 1 regardless of his type. On the other hand, if a sender does not receive a donation then his probability of survival is 1 − *a* if he is needy and 1 − *b* if he is healthy, where *a* > *b*. In addition, there is a relatedness parameter, *k* ∈ [0, 1], that captures the degree of common interest between the sender and the receiver–under any vector of strategies each player receives *k* times the payoff of the other player plus his or her own idiosyncratic payoff.

In the Sir Philip Sidney game, there is a breakdown in communication for certain regions of the parameters corresponding to a low cost of crying out, a low degree of relatedness, and a low cost of donation *d*. These are precisely the empirically relevant regions of costs that challenge the theory.

We introduce inattention into this environment quite simply: with some probability, *x*, the receiver observes the sender’s signal and does not with its complement (1 − *x*). We discover that no matter how small the signaling cost, as long it is strictly greater than zero,^2^ there is an interval of attention levels that sustains a separating equilibrium, in which (honest) communication occurs. Remarkably, some degree of inattention is always (weakly) optimal for the receiver: her maximal equilibrium payoff is no lower with partial inattention (*x* < 1) than with full attention (*x* = 1), and in certain regions of the parameter space the receiver is strictly better off when she is partially inattentive (her maximal equilibrium payoff is higher).

Next, we embed the signaling game into a larger game, in which the attention level, *x*, is an endogenous choice of the receiver made *ex-ante*, before the signaling game takes place.^3^ We discover that any subgame perfect equilibrium must yield the receiver a payoff at least as high as her maximal equilibrium payoff with full attention, and in certain parameter regions her equilibrium payoff is strictly higher.

When attention is endogenous, it is helpful in the following two ways. First, it provides a lower bound on the set of equilibrium payoffs of the game: since complete inattention may be chosen in the initial stage, at equilibrium the receiver can do no worse than the unique equilibrium payoff in the signaling game given complete inattention, where the two types of sender both remain silent. Second, and perhaps more compellingly, there is an interval of the attention parameter in which separating equilibria exist, despite the non-existence of such equilibria in the game with full attention. In short, inattentiveness enhances communication.

## 2 The Classic Sir Philip Sidney Game

We begin by revisiting the Sir Philip Sidney game (where we allow the signaling cost, *c*, to equal 0), which is depicted in Figure 1. There are two players, a sender (he) and a receiver (she); and the sender is one of two types, healthy or needy: Θ = {*θ*_*H*_, *θ*_*N*_}. The sender’s type is his private information, about which both sender and receiver share a common prior, *µ* := Pr (Θ = *θ*_*N*_).

**Figure 1:**
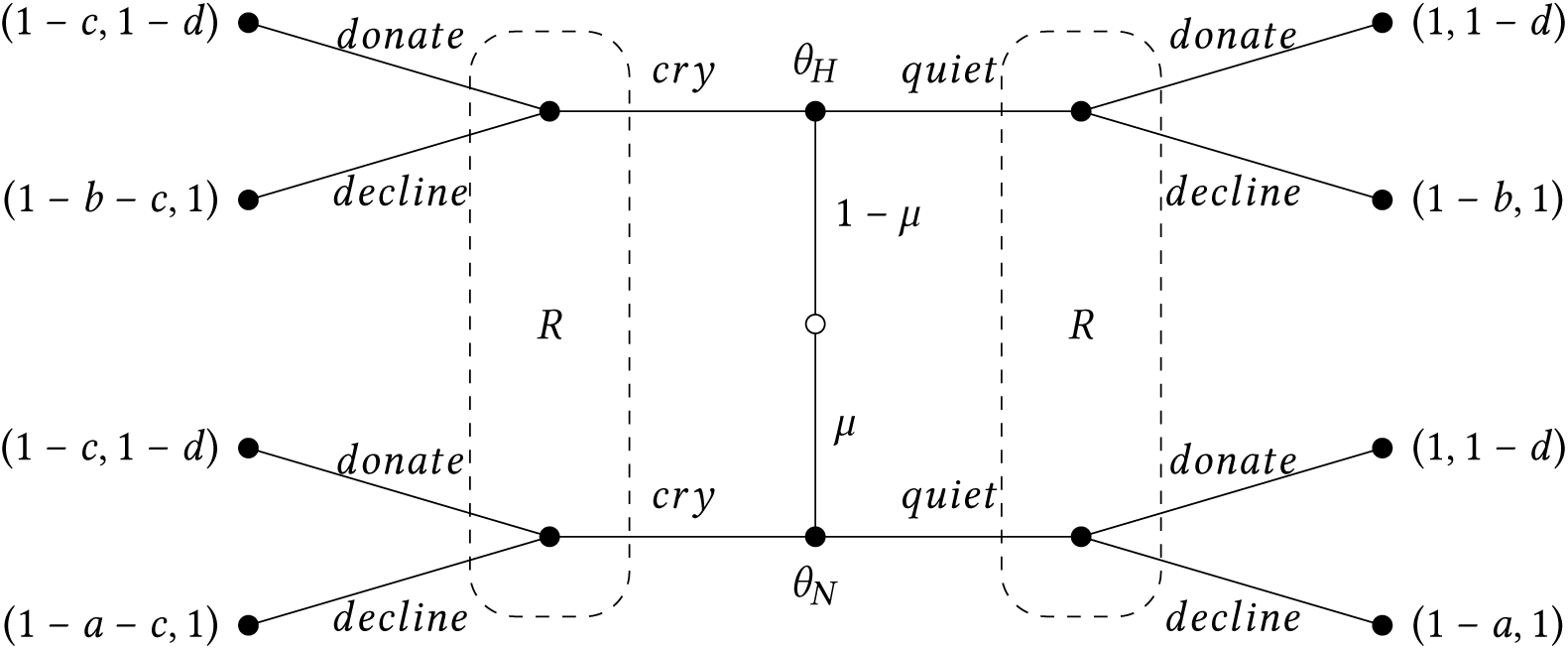
The Sir Philip Sidney Game

After being informed of his type, the sender chooses to either cry out (*cry*) or stay quiet (*quiet*). The receiver observes the sender’s choice of signal (but not his type), updates her belief about the sender’s type based on her prior belief and the equilibrium strategies and elects to either donate a resource (*donate*) or refuse to donate (*decline*). We impose that *a* > *b* and that *a, b, c*, and *d* take values in the interval [0, 1]. There is also a relatedness parameter *k* ∈ [0, 1]: after each outcome, a player receives his own payoff plus *k* times the payoff of the other player.

Throughout, we impose the following conditions:

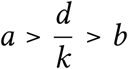

and

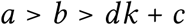

The first condition ensures that if the receiver is (sufficiently) confident that the sender is healthy then she strictly prefers not to donate and if she is sufficiently confident that the sender is needy then she strictly prefers to donate. The second condition eliminates any separating equilibria.

In addition, we define

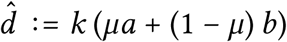

which saves us some room on the manuscript. Where convenient, we describe the equilibrium in the signaling game as a quadruple (·, ·; ·, ·), where the first entry corresponds to the strategy of *θ*_*H*_, the second entry to the strategy of *θ*_*N*_, the third entry to the response of the receiver to *quiet*, and the fourth entry to the response of the receiver to *cry*. In the case of pooling equilibria (equilibria in which both types of sender choose the same signal), we leave the response of the receiver to an off-path signal as · when there may be multiple responses that sustain an equilibrium.

The results of this section–which pertain to the standard discrete Sir Philip Sydney game– are standard in the literature. Naturally, they also correspond to special cases of the results with inattention. Consequently, all omitted proofs and derivations may be found in Appendix A (by setting the attention parameter, *x*, in each statement equal to 1). We begin with the following result, which follows from the parametric assumptions.

### Lemma 2.1.

*No separating equilibria exist*.

*Proof*. Standard, see e.g. Bergstrom and Lachmann (1997).

There do; however, exist pooling equilibria, both those in which both sender types choose *cry* and those in which both sender types choose *quiet*. Note that the pooling equilibrium in which both sender types choose *cry* requires that the receiver’s belief upon observing *quiet* (an off-path action) be such that she would at least (weakly) prefer to choose *decline* rather than *donate*. Moreover, the pooling equilibrium in which both sender types choose *quiet* and to which the receiver responds with *decline* also requires that the receiver’s belief upon observing *cry* (an off-path action) be such that she would at least (weakly) prefer to choose *decline* rather than *donate*. In some sense, this is less convincing of an equilibrium: shouldn’t the needy sender be more likely to cry out?^4^

The other pooling equilibrium, that in which both sender types choose *quiet* and the receiver responds with *donate*, makes no restrictions on the receiver’s off-path beliefs and is in that sense quite strong. Formally,

### Lemma 2.2.

*There exist pooling equilibria:*

1. (*cry, cry*; ·, *decline*) *is never an equilibrium;*
2. *If* 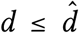 *then* (*cry, cry*; ·, *donate*) *is an equilibrium, and* (*quiet, quiet*; *donate*, ·) *is an equilibrium;*
3. *If* 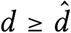, *then* (*quiet, quiet*; *decline*, ·) *is an equilibrium*.

*Proof*. Standard, see e.g. Bergstrom and Lachmann (1997).

There also exist equilibria in which players mix.^5^ Let *σ*_*i*_ denote the mixed strategy of a sender of type *θ*_*i*_:

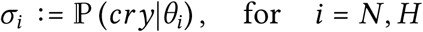

and let *λ* and *γ* denote the mixed strategies of the receiver following messages *cry* and *quiet*, respectively:

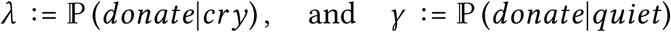

Then,

### Proposition 2.3.

*If* 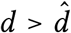, *there exists an equilibrium* (*σ*_*H*_, *cry*; *decline, λ*), *where*

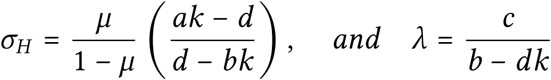

*If* 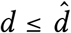, *there exist no equilibria in which type θ*_*H*_ *mixes and type θ*_*N*_ *chooses cry*.

Similarly,

### Proposition 2.4.

*If* 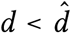, *there exists an equilibrium* (*quiet, σ*_*N*_; *γ, donate*), *where*

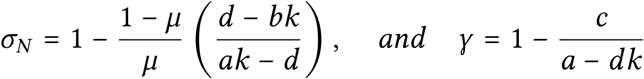

*If* 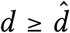, *there exist no equilibria in which type θ*_*N*_ *mixes and type θ*_*H*_ *chooses quiet*.

At first glance, these equilibria might seem better for the receiver than the pooling equilibria. Indeed, at least some information is transmitted in them, whereas when the senders pool there is no information transmission whatsoever. However, in these mixed strategy equilibria the information conveyed is, in a sense, useless. Indeed, it is easy to see that there cannot be any equilibria in which useful information is transmitted. That is, there can be no equilibria in which different actions are strictly preferred after different messages, since then there would be a sender type who could deviate profitably to the message that is followed by *donate*. Thus, all equilibria are ones in which the receiver is (weakly) willing to do the same action following any message. Of those equilibria, the best for the receiver is the one in which the sender types pool on *quiet*.

Accordingly, if 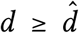 then the receiver optimal equilibrium is the pooling equilibrium (*quiet, quiet*; *decline*, ·), which yields her a payoff of

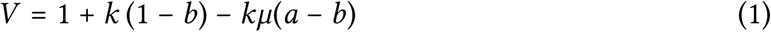

and if 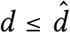 then the receiver optimal equilibrium is the pooling equilibrium (*quiet, quiet*; *donate*, ·), which yields her a payoff of

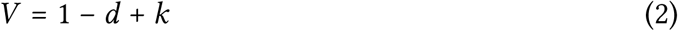

There exist no other equilibria for parameters in the regions that we specified. Indeed,

### Proposition 2.5.

*There exist no equilibria in which*

1. *Type θ*_*H*_ *mixes and type θ*_*N*_ *chooses quiet; or*
2. *Type θ*_*N*_ *mixes and type θ*_*H*_ *chooses cry*.

This proposition is easy to comprehend. First, if the healthy sender mixes and the needy sender stays quiet, then after observing *cry* the receiver is sure that the sender is healthy and so does not donate. This is the worst combination for the sender (incur the cost of *cry* yet not receive a donation) and so the healthy type strictly prefers to remain quiet. Second, if the needy sender mixes and the healthy sender cries, then after observing *quiet* the receiver is sure that the sender is healthy and so donates. This, on the other hand, is the best combination for the sender (he does not incur the crying cost yet receives a donation) and so again the healthy type strictly prefers not to cry out.

### 2.1 The Value of Information

Note that we may rewrite the condition 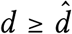 as

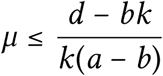

and so we can write the receiver-optimal payoff as a function of the belief

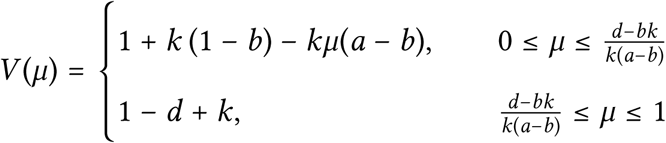

Evidently, *V* is convex in *µ*. An interesting ramification of this is that any (free) *ex-ante* information about the sender’s type benefits the receiver. This is not generally true in communication games even if there are just two types and two receiver actions (when the receiver’s payoff is directly affected by the sender’s message).^6^ However, it is true here: any free information benefits the receiver.

The receiver’s optimal equilibrium payoff is depicted in Figure 2 for the following values of the parameters: *a* = 1, *b* = 3/8, *c* = 11/64, *d* = 1/8, and *k* = 1/4.

**Figure 2:**
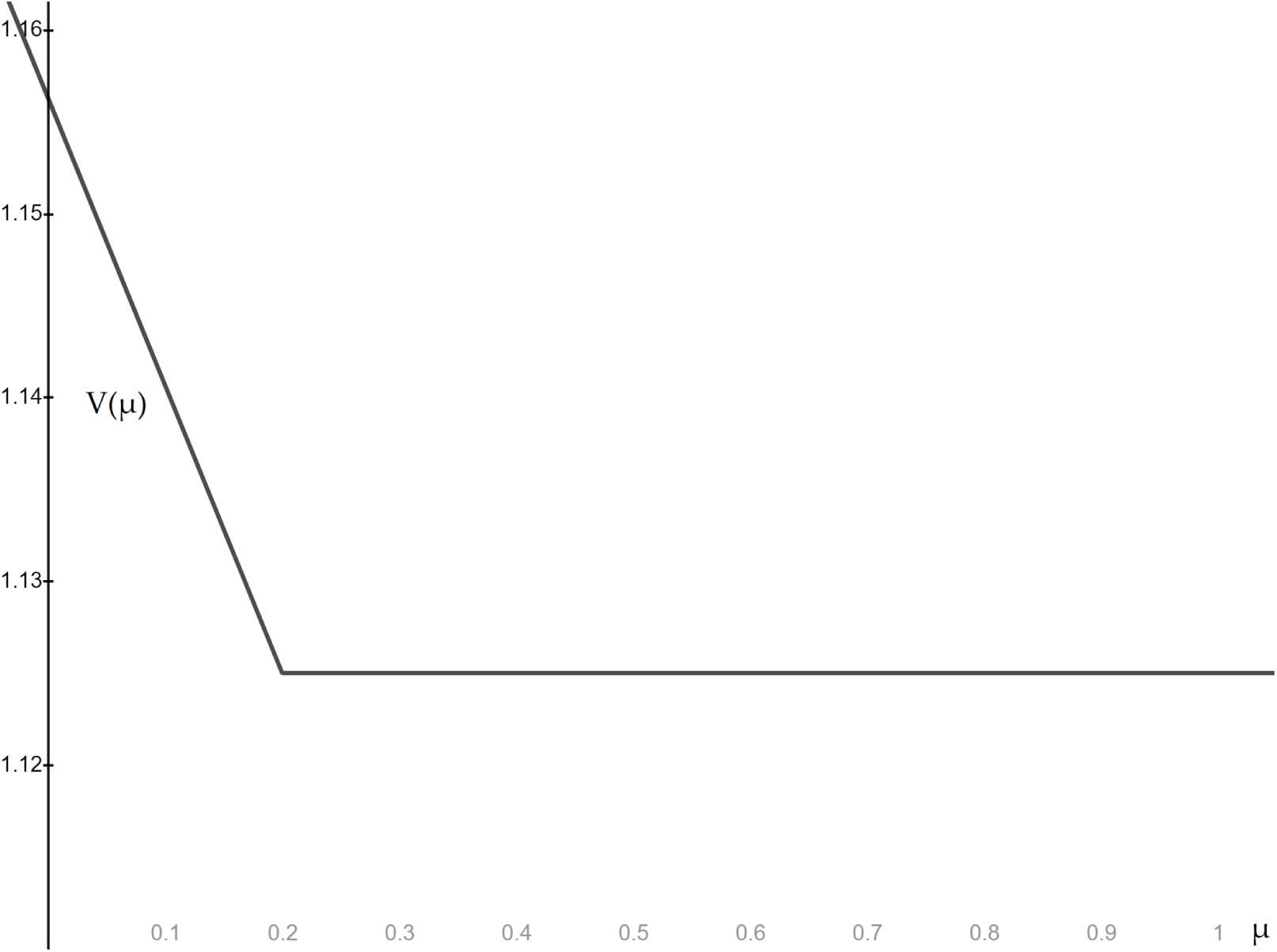
Receiver’s Payoff, No Inattention

## 3 Inattention

Now let us explore the notion that full attention may not be generally optimal for the receiver. We modify the game by assuming that the receiver is inattentive–she observes the sender’s signal with some probability *x* and does not with its compliment. Introducing inattention requires that we make a modelling decision in response to this question: does the receiver know when she is inattentive? That is, can she distinguish between the absence of a signal due to inattention and *quiet*?

We introduce the following terms:

### Definition 3.1.

The receiver is **Consciously Inattentive** if she is aware when she is inattentive. That is, she can distinguish between *quiet* and the absence of a signal due to inattention (Ø).

Conversely,

### Definition 3.2.

The receiver is **Unconsciously Inattentive** if she is unaware when she is inattentive. That is, when she observes *quiet*, she does not know whether it was due to her inattention or *because there was no signal to observe*.

For the majority of this paper we assume that the receiver *can* distinguish between the absence of a signal due to inattention and *quiet*–she is consciously rather than unconsciously inattentive. As we subsequently show, in this game, conscious inattention is always weakly better for the receiver than unconscious inattention. Furthermore, we briefly explore unconscious inattention in Section 4 and argue there that our findings are qualitatively unaffected by the type of inattention.

Conscious and Unconscious Inattention are illustrated in Figure 3. With conscious inattention, if the receiver does not attend to the signal then she simply makes the optimal choice given her information at hand, which is merely her prior. Hence, she chooses *decline* if and only if 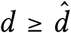 and *donate* otherwise. With unconscious inattention the receiver observes either *quiet* or *cry*, and must account for her inattention when she chooses her best response following *quiet*.

**Figure 3:**
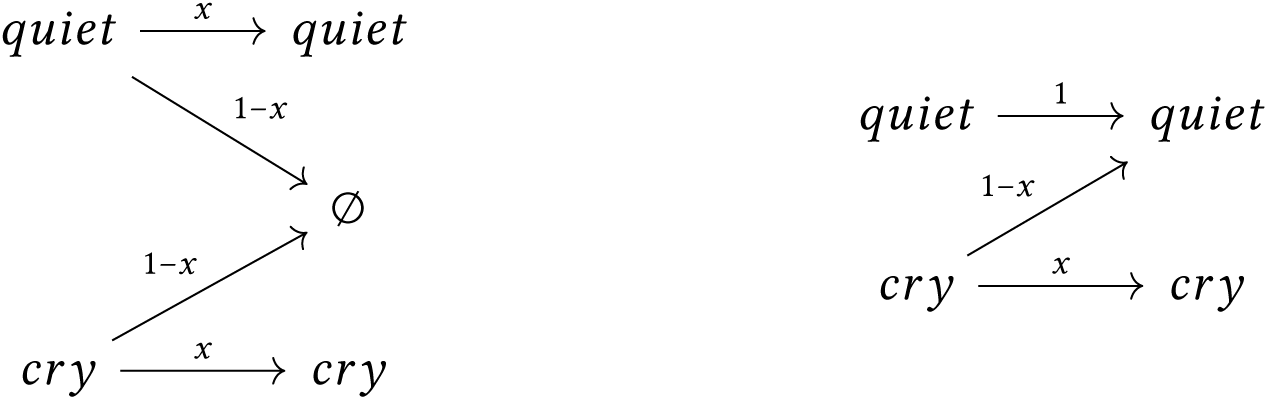
Conscious and Unconscious Inattention

Focusing on the conscious attention case, we see that there are two critical cutoff beliefs of *x*, 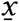 and 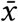, which divide the range of possible values of *x* into three regions, depicted in Figure 4. Explicitly,

**Figure 4:**
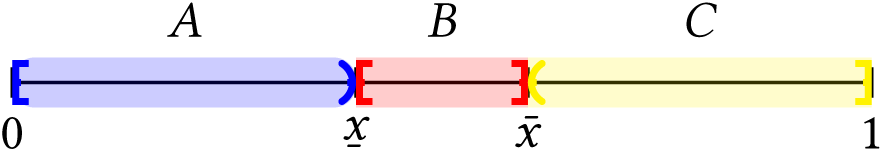
The Critical Regions of *x*

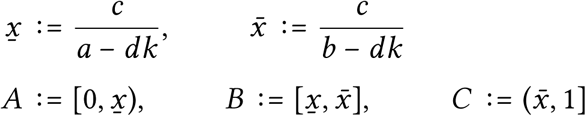

The first result highlights that, in contrast to the first section of this paper, in which there did not exist separating equilibria, other values of *x* may beget separation.

### Lemma 3.3.

*If x* ∈ *B then there exist separating equilibria in which θ*_*H*_ *chooses quiet and θ*_*N*_ *chooses cry. Such equilibria yield the following payoffs to the receiver:*

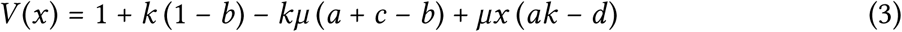

*when* 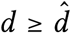, *and*

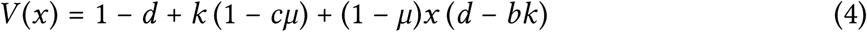

*when* 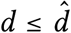. *The level of attention that maximizes the receiver’s payoff is*

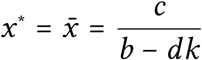

*Proof*. First, let 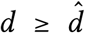, so that, when attending, *decline* is the receiver’s response to not observing. The receiver’s payoff reduces to

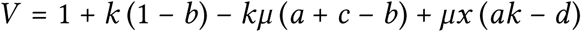

which is increasing in *x*. Type *θ*_*H*_ ‘s incentive constraint reduces to

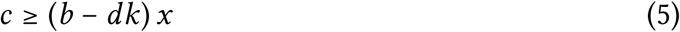

and type *θ*_*N*_ ‘s incentive constraint reduces to

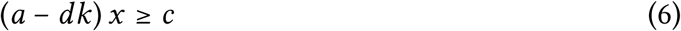

Thus, a separating equilibrium of this form exists provided the attention parameter, *x*, satisfies

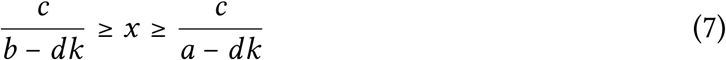

Since *V* is obviously increasing in *x*, the value of the attention parameter that maximizes the receiver’s payoff is

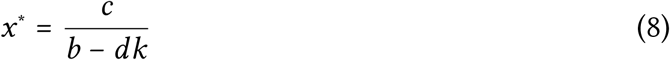

When 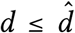, the same procedure suffices *mutatis mutandis*. Any value of the attention parameter, *x*, that satisfies the inequalities in Expression 7 begets a separating equilibrium, and the value of the attention parameter, *x*^∗^, that maximizes the receiver’s payoff is given in Equation 8.

The intuition behind this result is simple. If the receiver is moderately inattentive (*x* ∈ *B*), the receiver can lessen the incentive of either type to deviate and mimic the other. Consequently, Inequalities 5 and 6 are key, which illustrate how moderate inattention rescales the two sender types’ marginal benefits and engenders separation. Then, the receiver-optimal value of the attention parameter, *x*^∗^, is the value that leaves type *θ*_*H*_ indifferent between separating and deviating to mimic *θ*_*N*_. The receiver would like to pay the maximal amount of attention such that she is sufficiently inattentive for the sender types to separate.

It is easy to see that no matter the attention parameter, *x*, the “opposite” separating equilibrium cannot exist:

### Lemma 3.4.

*There exists no attention parameter x* ∈ [0, 1] *such that there exists a separating equilibrium in which θ*_*H*_ *chooses cry and θ*_*N*_ *chooses quiet*.

Of course, there also exist pooling equilibria. *Viz*,

### Lemma 3.5.

*There does not an exist an x* ∈ [0, 1] *such that* (*cry, cry*; ·, *decline*) *is an equilibrium. Moreover*,

1. *For* 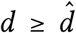, (*quiet, quiet*; *decline*, ·) *is an equilibrium. The receiver’s resulting payoff is given in Expression* 1.
2. *For* 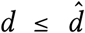, (*quiet, quiet*; *donate*, ·) *is an equilibrium*. (*cry, cry*; ·, *donate*) *is also an equilibrium if and only if* 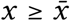. *The receiver’s resulting payoff is given in Expression* 2.

The receiver-optimal pooling equilibria are those in which the sender types remain silent, and those equilibria are optimal (among all equilibria pooling or otherwise) for all *x* ∈ *A* and *x* ∈ *C*. Note that there is only an equilibrium in which the sender types both choose *cry* if both *d* and *x* are sufficiently high. In particular, for any *d*, if *x* is sufficiently low then there is no equilibrium in which the sender types both *cry*.

Next, the following lemma establishes that if *x* ∈ *B*, it is possible that the separating equilibrium is optimal. To wit,

### Lemma 3.6.

*Let* 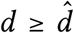 *and x* ∈ *B. Then the equilibrium that maximizes the receiver’s payoff is* (*quiet, cry*; *decline, donate*) *if and only if*

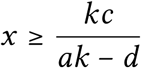

*Otherwise it is* (*quiet, quiet*; *decline*, ·).

*Let* 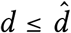 *and x* ∈ *B. Then the equilibrium that maximizes the receiver’s payoff is* (*quiet, cry*; *decline, donate*) *if and only if*

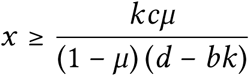

*Otherwise it is* (*quiet, quiet*; *donate*, ·).

*Proof*. First, let 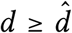. Using the receiver’s payoff from the pooling equilibrium (Expression 1) and her payoff from the separating equilibrium (Expression 3), we have *V*^*sep*^(*x*) ≥ *V*^*pool*^ if and only if

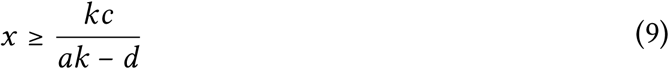

Note also that if *x* = *x*^∗^ then this becomes

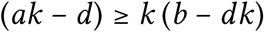

Second, let 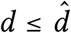. Then, *V*^*sep*^(*x*) ≥ *V*^*pool*^ if and only if

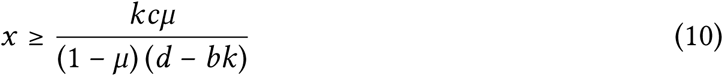

If *x* = *x*^∗^ then this becomes

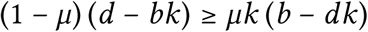

Pausing briefly to look at the cutoffs above which separation is better, we see that the right-hand side of Inequality 9 is increasing in both *c* and *d* and decreasing in *k* and *a*. Hence, small^7^ decreases in the signaling and/or donation costs enlarge the set of attention parameters *x* such that separation is better for the receiver than pooling. Analogously, small increases in the relatedness parameter and/or the cost suffered by the needy type have the same effect.

The right-hand side of Inequality 10 is also increasing in *c* and decreasing in *k*, and so the same logic holds. However, it is now increasing in *b* and in *µ* and decreasing in *d*. It is easy to see why it should be increasing in *µ*: as the proportion of needy types increases, the uninformativeness of pooling is not as harmful to the receiver (recall that since 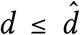 the receiver is donating). Similar reasoning explains the relationship with *d*: as *d* increases, pooling becomes more costly since donation itself is more costly. As *b* increases it becomes harder to induce separation (the healthy type has a stronger incentive to mimic the needy type) which reduces the receiver’s benefit from the separating equilibrium.

Let us also revisit *x*^∗^ and the properties of the optimal attention level. Directly, we have

### Lemma 3.7.

*x*^∗^ *is decreasing in b and increasing in c, d, and k*.

This result agrees with our intuition: as signaling costs, donation costs, and relatedness increase, so does the amount of attention paid by the receiver at the optimum. This optimal attention level decreases, on the other hand, as the healthy type’s loss from not receiving a donation increases. Thus, an increase in a parameter that helps drive communication (separation), or a decrease in the parameter that helps thwart communication, increases the optimal level of attention.

There also exist mixed strategy equilibria under limited attention. Recall that *σ*_*i*_ denotes the probability that a sender of type *θ*_*i*_ chooses *cry*; and that *λ* and *γ* denote the probabilities that the receiver chooses *donate* following messages *cry* and *quiet*, respectively.

The first proposition establishes that there are a variety of equilibria in which the healthy type of sender mixes and the needy type cries.

### Proposition 3.8.

*If* 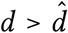 *and*

1. 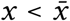, *there exist no equilibria in which θ*_*H*_ *mixes and θ*_*N*_ *chooses cry;*
2. 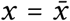, *there exists a continuum of equilibria* (*σ*_*H*_, *cry*; *decline, donate*), *where*

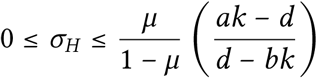
3. 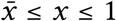, *there exists an equilibrium* (*σ*_*H*_, *cry*; *decline, λ*), *where*

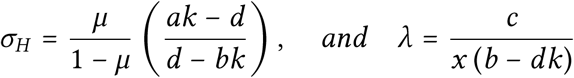

*If* 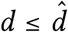 *there exists an equilibrium in which θ*_*H*_ *mixes and θ*_*N*_ *chooses cry if and only if* 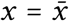.

The second proposition establishes that there are also a variety of equilibria in which the needy type of sender mixes and the healthy type does not cry.

### Proposition 3.9.

*If* 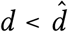 *and*

1. 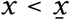, *there exist no equilibria in which θ*_*N*_ *mixes and θ*_*H*_ *chooses quiet*.
2. 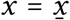, *there exists a continuum of equilibria* (*quiet, σ*_*N*_; *decline, donate*), *where*

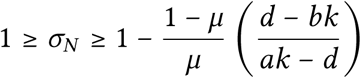
3. 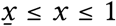, *there exists an equilibrium* (*quiet, σ*_*N*_; *γ, donate*), *where*

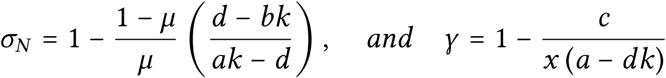

*If* 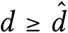 *there exists an equilibrium in which θ*_*N*_ *mixes and θ*_*H*_ *chooses quiet if and only if* 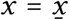.

On the other hand, even limited attention cannot sustain some mixed strategy equilibria. Namely,

### Proposition 3.10.

*There exist no equilibria in which*

1. *Type θ*_*H*_ *mixes and type θ*_*N*_ *chooses quiet; or*
2. *Type θ*_*N*_ *mixes and type θ*_*H*_ *chooses cry*.

Just like the case with full attention, there can be no equilibria where the healthy sender fully identifies himself by choosing *cry* (the first case), or where the needy sender fully identifies himself by choosing *quiet* (the second case).

### 3.1 Empirical Relevance

Before proceeding onward, we make a brief digression to assure ourselves that limited attention is realistic. It is: not only are animals inattentive^8^ but there is considerable empirical evidence that inattention affects their behavior in significant ways. One well-established result is that foragers searching for food (or fighting among themselves) may be less likely to notice approaching predators when they are intently focused. This effect has been documented in a variety of animals including sticklebacks (Milinski (1984)), guppies (Godin and Smith (1988) and Krause and Godin (1996)), blue jays (Dukas and Kamil (2000) and Dukas and Kamil (2001)), blue tits (Kaby and Lind (2003)), and cichlids (Ota (2018)). Similarly, Chan et al. (2010) find that (anthropogenic) sounds may occupy the attention of hermit crabs, leaving them more vulnerable to predation. Dukas (2002) writes, “Animals must commonly handle more than one task at a time. An ubiquitous example is searching for food while avoiding predators.” Obviously, a similar dual concern would exist in the sort of parent-offspring interaction analyzed in this paper.

There is also evidence that limited attention plays a role in courtship. Dukas (2002) writes, “The fact that an animal has the sensory capacity to perceive certain information does not necessarily imply that it actually attends to that information while…watching courtship displays.” Richards (1981) states that “a great many” birds have calls structured so that they are initially very detectable (and notes that “a similar structure exists in the long-range calls of some primates”). Multiple studies have found related attention-grabbing behavior in anoles; see, e.g., Ord and Stamps (2008) and Ord (2012).

Such evidence informs the theoretical model developed by Számadó (2015),^9^ who modifies a basic signaling game by embedding it into a larger interaction in which search (by the receiver) and attention-seeking behavior (by the sender) play vital roles. In his model, sender and receiver are initially far apart and the receiver must search (at a cost) in order to find the sender. This search can be made easier by the sender, who may, at the beginning of the interaction, give a (costly) attention seeking display, which makes it more likely that the receiver will find the sender. Once the receiver has found the sender, a standard signaling game of resource donation takes place.

Számadó shows that there can be equilibria in which giving attention-seeking displays can be part of a subgame perfect equilibrium (and part of an evolutionarily stable strategy) in the game. Note that unless the signaling cost for the healthy type is 0 or negative (i.e. the signal is intrinsically *beneficial*) the additional stages cannot beget separation in the signaling portion of the game when such an honest equilibrium does not exist in game absent the display and search stages. However, the extra stages can allow for separation at the early stage, when the displays are given, and so that sort of honesty can manifest.

It seems natural to think of inattention in the context of Számadó’s work. The introduction of the additional stages notwithstanding, there remain regions of the parameter universe in which no separation at any stage is possible. This transpires (or rather separation cannot transpire) if the cost of sending a display is too low and/or the likelihood of the receiver finding the sender without a display is too high, in conjunction with the parameters in the signaling subgame being such that there is no separation in the subgame. Accordingly, the analysis above can be seen as complementary to Számadó (2015), since inattention would play an analogous role when applied to Számadó’s paradigm. In particular, we could think about the benefits to the receiver of inattentiveness towards the sender’s *display*.

### 3.2 What if the Receiver Could *Choose* Her Level of Attention?

Until now, we have treated inattention–or more specifically the attention level, *x*–as an exogenous primitive of the game. Let us briefly explore the ramifications of treating it as a choice variable. Namely, now let there be an initial stage in which the receiver *chooses* and *publicly*^10^ commits to her level of attention, *x*. Following this choice, the signaling game proceeds in the standard manner under parameter *x*.

In the first result, we discover that when *x* is endogenous, the set of Nash equilibria of the game is quite large.

#### Theorem 3.11.

*Let E*(*x*) *be a Nash equilibrium of the signaling subgame with an exogenously given x and ε*(*x*) *be the set of all such equilibria. Then for all E*(*x*) ∈ *ε* (*x*), *for all x* ∈ [0, 1], *there exists a Nash equilibrium of the game in which attention level x is chosen and E*(*x*) *is played in the signaling subgame*.

*There are no Nash equilibria of the game in which, following the on-path choice of attention parameter, x, the vector of strategies for the signaling subgame is not a Nash equilibrium of the subgame*.

That is, any equilibrium in any signaling subgame that follows an on-path choice of parameter *x* is part of a Nash equilibrium of the game, and the vector of strategies in any on-path signaling subgame must constitute an equilibrium of the subgame.

*Proof*. The proof of this result is nearly trivial. Indeed, the first part of the theorem’s statement requires only that we construct strategies that prevent the receiver from deviating to some other *x*′not equal to the on-the-equilibrium-path *x*. That is easy–we merely have the sender types pool on *cry* following any deviation. This may be an incredible threat, but since this is a Nash equilibrium and not a subgame perfect equilibrium that is fine.

The second part of the statement is obvious–by definition, if the vector of strategies of the on-path signaling subgame is not an equilibrium of the subgame then at least one player has a profitable deviation.

Second, we look for subgame perfect equilibria. We will use the following conditions in the statement of the results:

#### Condition 3.12.

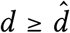 and Inequality 9 holds.

#### Condition 3.13.

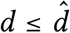 and Inequality 10 holds.

Then,

#### Theorem 3.14.

*Suppose there exists some* 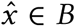 *such that either Condition 3*.*12 or Condition 3*.*13 holds. Then, there exists a collection of subgame perfect equilibria consisting of a choice of* 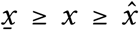 *in the first stage, and* (*quiet, cry*; *decline, donate*) *in the signaling portion of the game. The receiver optimal subgame perfect equilibria have x* = *x*^∗^ = *c*/(*b* − *dk*).

*Proof*. This follows from Lemma 3.6. Subgame perfection is obtained since the sender types are willing to pool following any deviation by the receiver.

Next, we see that there are always subgame perfect equilibria in which no information is transmitted.

#### Theorem 3.15.

*There is always a collection of subgame perfect equilibria consisting of any choice of x in the first stage, and* (*quiet, quiet*; *decline*, ·) *for* 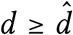, *and* (*quiet, quiet*; *donate*, ·) *for* 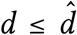 *in the signaling portion. If neither Condition 3*.*12 nor Condition 3*.*13 holds then this collection is unique*.

*Proof*. This follows from the fact that the equilibria in which both sender types are *quiet* are the pooling equilibria that (uniquely) maximize the receiver’s payoff. Since these are the unique equilibrium should the receiver choose any *x* ∈ *A*, they must be the equilibria played for any *x* since otherwise the receiver would have a profitable deviation in the initial stage to an *x* ∈ *A*.

This pair of theorems illustrates the two main effects of allowing the receiver to choose her level of attention initially. First, limited attention yields separating equilibria even when such equilibria could not exist under full attention. That is, honest communication arises in a scenario in which the conflict between the receiver and the sender would typically be too great for it to occur. Second, enabling the receiver to choose her level of attention ensures that the equilibrium played in the signaling portion of the game is relatively “good” for the receiver (either best or second-best) and provides a lower bound for the receiver’s payoff.

#### 3.2.1 A Comment on Public Commitment

It is important to keep in mind that when the choice of attention level, *x*, is endogenous we require that this choice be public. We can dispense with this assumption, but only if we modify the game. Why is this necessary? Well, suppose that *x* were not public but instead a private choice of the receiver. If we maintained the assumption that information was free then any Subgame Perfect Equilibrium would be one in which the sender types pooled on quiet. This is due to the incentive for the receiver to “sneak a peak” and secretly deviate to full attention.

However, there are a few natural modifications or extensions that allow for a private choice of attention level:

1. **Costly Attention:** We could impose that it is costly to be attentive, i.e. that the receiver incurs some attention cost *w*(*x*) that is increasing (and twice continuously differentiable and convex, for convenience) in *x*. This is quite natural–there is an opportunity cost to attention.^11^ A bird does not want to devote all of her attention to her chicks because that would leave her too vulnerable to a predator. A predator does not want to focus exclusively on the stotting behavior of the gazelle for similar reasons. In a separating equilibrium, the receiver’s payoff would now b

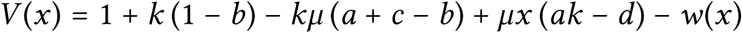

and taking the first order conditions we see that the attention level 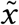 that solves

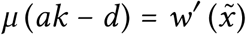

begets such a separating equilibrium provided 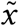.
2. **Reputation:** We could modify the scenario to a long-run interaction between a long-lived receiver and a sequence of short-lived (or very impatient) senders. Provided there was a small possibility that the receiver was committed to the optimal attention level, she could ensure herself an average payoff almost as high as her optimal inattention payoff in every equilibrium. The argument for this statement proceeds as follows. Theorem 3.16, *infra*, establishes that inattention is as good as the receiver-optimal medium; Proposition 3.5 in Whit-meyer (2019a) shows that the receiver-optimal medium is as good as commitment; and Theorem 3.1 in Fudenberg and Levine (1992) states that a sufficiently patient long-run player obtains a payoff virtually as high as her commitment payoff in any equilibrium.
3. **Multiple Types of Receiver:** Another solution is to impose that the population of receivers consists of two types, those who are fully attentive and those who are completely inattentive. It is clear that if the likelihood that a receiver is attentive falls within the interval *B* then there is an equilibrium in which the sender types signal honestly.

In addition, public commitment to inattention may not be as implausible as it initially seems. A receiver could deliberately engage in a distracting task that she knows will make it difficult for her to observe a signal. This behavior is quite sophisticated, but anecdotally it seems like people, at least, do behave in this way. For instance, think of a teenager in her room who deliberately puts headphones on so that her parents’ questions may not be heard.

### 3.3 Inattention Corresponds to the Receiver-Optimal Medium

A paper close in spirit to this one is Lachmann and Bergstrom (1998), who allow for perceptual error on the part of the receiver and introduce the notion of a *medium*, which distorts the signals observed by the receiver.^12^

In the analysis above we restricted the set of media the receiver can choose to those of a specific sort, those which correspond to inattention, and then endogenized the medium by making it a choice of the receiver. Remarkably, as we discover in Theorem 3.16, the receiver-optimal equilibrium under her optimal choice of attention remains supreme even were she able to choose any medium, however complex.

#### Theorem 3.16.

*V*^*med*^ *be the receiver’s payoff for the receiver-optimal equilibrium under the best-possible medium for the receiver. Then there exists an attention parameter x such that V* (*x*) = *V*^*med*^.

*If either Condition 3*.*12 or Condition 3*.*13 holds then the optimal parameter is x*^∗^ = *c*/(*b* − *dk*). *If neither holds then any parameter x* ∈ [0, 1] *is optimal*.

*Proof*. We wish to choose a medium in order to maximize *V*, which we will then show coincides with the receiver’s payoff under inattention. From Whitmeyer (2019a) it is without loss of generality to restrict our attention to pure strategies of the sender types. Moreover, Theorem 3.8 in Whitmeyer (2019a) establishes that we need only consider a (relaxed) *commitment* problem for the receiver. That is, suppose that the receiver can commit to choosing *donate* with probability *p* and *decline* with probability 1 − *p* following *cry*; and *donate* with probability *q* and *decline* with probability 1 − *q* following *quiet*. The receiver solves the following optimization problem,

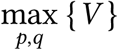

subject to

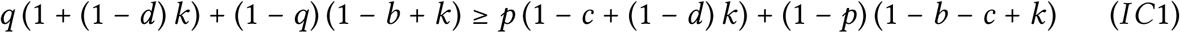

and

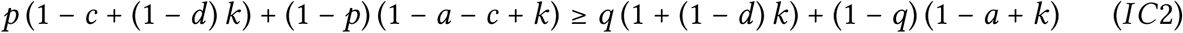

where

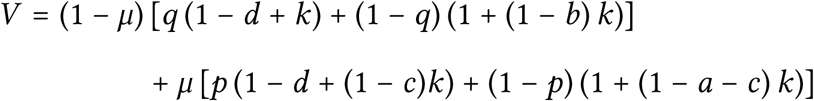

This optimization problem is easy to solve and yields *q* = 0 and *p* = *c*/(*b* − *dk*) for 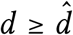, and *q* = 1 − *c*/(*b* − *dk*) and *p* = 1 for 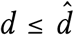. Substituting these into the value function, we obtain *V* (*x*^∗^). If either Condition 3.12 or Condition 3.13 holds, then this maximizes the receiver’s payoff, and as illustrated in Whitmeyer (2019a), since *p* and *q* solve the commitment problem, this must be the solution to the problem of choosing an optimal medium.^13^

On the other hand, if neither condition holds then the result is trivial. The receiver-optimal equilibrium is one in which both types pool on *quiet* and is attainable under any *x*.

### 3.4 The Value of Information with Optimal Attention

As we did earlier, the receiver’s payoff can be written as a function of the prior, *V*_*I*_ (where *I* stands for inattention). From Whitmeyer (2019a), the receiver’s payoff when she may choose the optimal medium is convex in *µ*. Thus, here, since inattention corresponds to the optimal medium, *V*_*I*_ must be convex. That is, any (free) *ex-ante* information about the sender’s type must benefit the receiver at least weakly. Substituting the parameter values used *supra*^14^, the receiver’s payoff function is

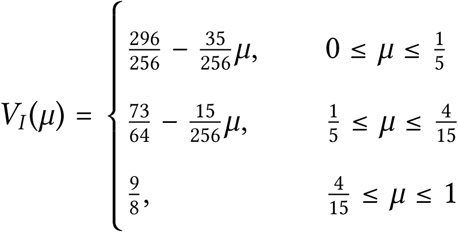

This function is depicted in Figure 5. The receiver’s payoff without inattention (from Figure 2) is the dashed black curve and her payoff with optimal inattention is the green curve.

**Figure 5:**
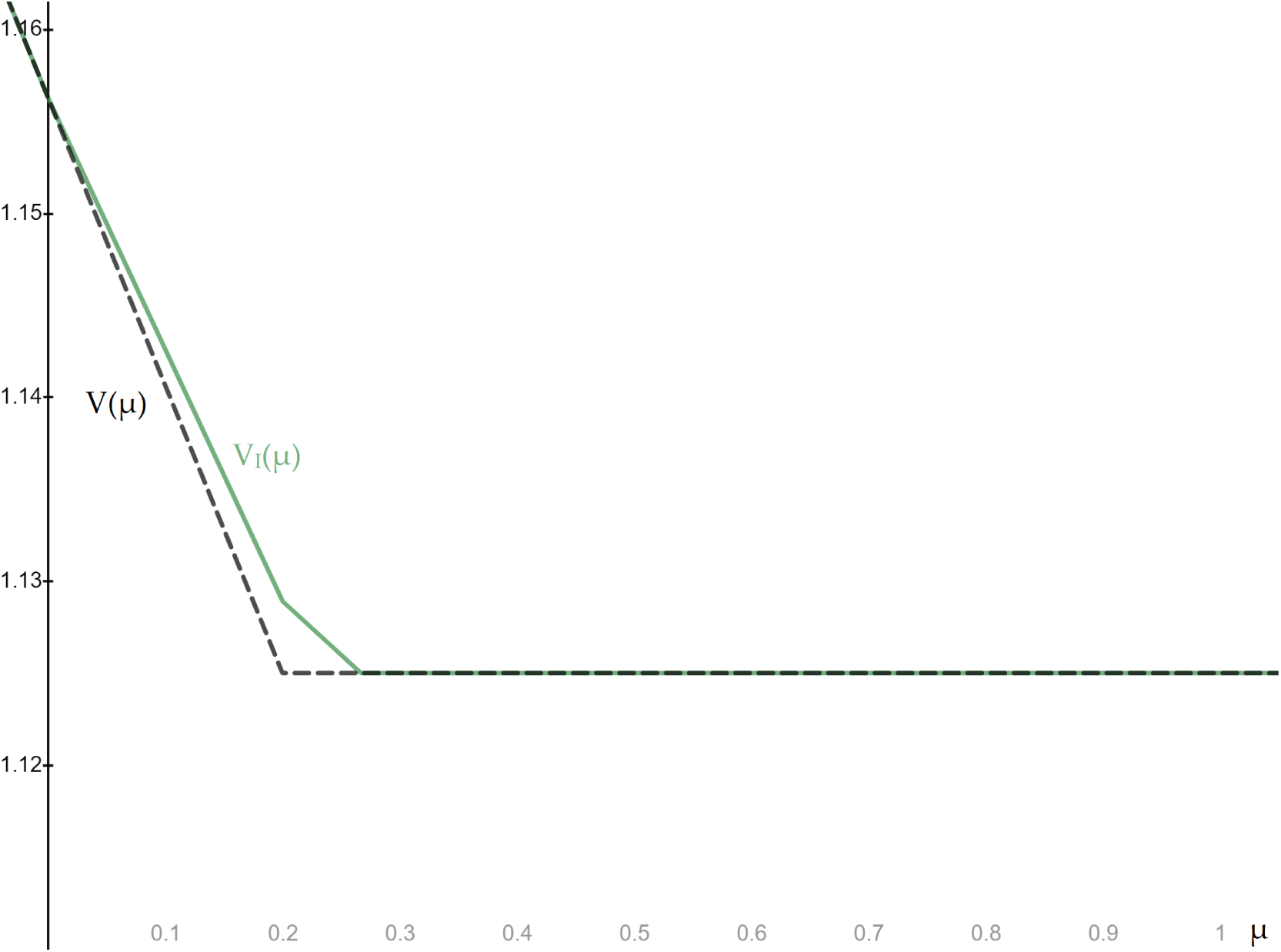
Receiver’s Payoff, Inattention

## 4 Unconscious Inattention

Here, we briefly explore the case of unconscious inattention. We establish that an analog to Lemma 3.3 holds: unconscious inattention can also beget separation. This comes with something of a *caveat*; however, we go to note that conscious inattention is always weakly better for the receiver than unconscious inattention. We finish this section by considering an extension in which the sender can affect the fidelity of the medium through his signal.^15^ He may choose the volume of his cry, and the louder the cry the more likely the receiver is to observe it. Again, we discover an analog to Lemma 3.3.

Let us begin in the standard framework of the paper but impose that the receiver is unconsciously inattentive.

### Lemma 4.1.

*If the attention parameter x* ∈ *B and*

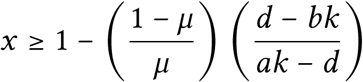

*then there exists a separating equilibrium in which θ*_*H*_ *chooses quiet and θ*_*N*_ *chooses cry. A stronger sufficient condition for such an equilibrium is x* ∈ *B and* 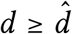. *The receiver’s payoff is given in Equation* 3. *The level of attention that maximizes the receiver’s payoff is*

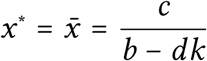

*Proof*. The incentive constraints for the sender types are the same as in Lemma 3, i.e., *x* must lie in the interval *B*. Moreover, the receiver’s decision must also be sequentially rational, and so, using Bayes’ law, this reduces to

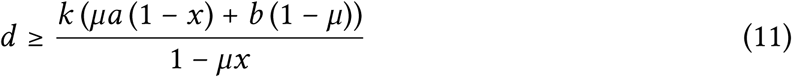

Observe that the right hand side of this inequality is decreasing in *x* and that when *x* = 0 the inequality simplifies to just 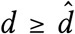. Thus, a sufficient condition for the existence of such an equilibrium is that *x* ∈ *B* and 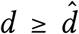. Obviously, *x* ∈ *B* is also a necessary condition (or else a sender type would have a profitable deviation). Rearranging Inequality 11 yields the inequality given in the proof’s statement.

The remaining equilibria are qualitatively similar to the scenario with conscious inattention (and indeed to the full attention setting). The same pooling equilibria exist, and there exist similar mixed strategy equilibria, with slight modifications to the precise mixed strategies that support the equilibria. As in the case with conscious inattention, there exist no equilibria where the healthy type mixes and the needy type stays quiet or where the needy type mixes and the healthy type cries.

Naturally, Theorems 3.14 and 3.15 hold as well: with unconscious inattention the set of subgame perfect equilibria consists of arbitrary choices of *x* followed by pooling in the signaling subgame and, under certain parametric conditions, choices of *x* that sustain separation followed by separation in the signaling subgame.

The proof of Lemma 4.1 hints at the next result. Observe that in contrast to the conscious inattention paradigm, there is an additional constraint, which will not be satisfied if the receiver is too confident that the sender is type *θ*_*N*_ (if *µ* is too high). Since Theorem 3.16 implies that the optimal level of (conscious) inattention for the receiver *always* allows her to receive a payoff as high as through the optimal medium, we have

### Corollary 4.2.

*The receiver’s maximal payoff with conscious inattention is* (*weakly*) *higher that her maximal payoff with unconscious inattention*.

The game with exogenous unconscious inattention is quite similar to the “Pygmalion game” as introduced by Huttegger et al. (2015).^16^ There, the receiver only observes a sender’s signal with some type-dependent probability, and when she makes her decision she is in just one of two possible information sets: she knows whether she has observed a signal, but does not know whether a signal was sent when no signal is received. The existence of an honest signaling equilibrium in the Pygmalion game depends on both the signaling cost and the likelihood of transmission success. As in this paper, success probabilities less than 1 can encourage honesty, and allowing for heterogeneity in such probabilities can provide an even stronger impetus if the signals of high types (which correspond to the needy types in this paper) are more likely to be observed.

I have noted that this work complements Számadó (2015); it also complements Huttegger et al. (2015). In Huttegger et al. (2015), if the cost of signaling is too low and the likelihood of a signal being observed is too high, then, just like the basic Sir Philip Sidney game, there are no honest equilibria. Thus, there is room for inattention on the part of the receiver, which would lower the likelihood of a successful signal and thereby engender separation (honest behavior by the sender types).

### 4.1 Piercing the Silence

Now let us modify the setting by allowing the sender’s signal to alter the fidelity of the medium. We make the following change to the basic game: instead of choosing *cry* or *quiet* the sender chooses how loud to make her cry. He chooses any 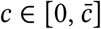, where *c* is both the volume of the cry as well as the cost incurred by a cry of such volume. Simply, the louder the cry, the more likely a predator is to hear it.^17^

The remaining parameteric assumptions are unchanged, and we assume that

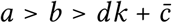

which eliminates any separating equilibria. Again, the receiver optimal equilibrium without inattention is one where both types of sender choose *c* = 0, i.e., they remain silent.

Now suppose that the receiver is unconsciously inattentive. The louder the sender’s cry the more likely it is to break the receiver’s *reverie*. Formally, given a cry of volume *c*, the receiver observes (hears) the cry with probability *x*(*c*), where

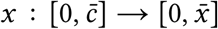

is a concave, strictly increasing, twice continuously differentiable function. Moreover, *x*(0) = 0 and because *x*(*c*) is a probability, 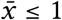.

#### Proposition 4.3.

*For any x*(·) *and c* ∈ 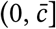 *such that*

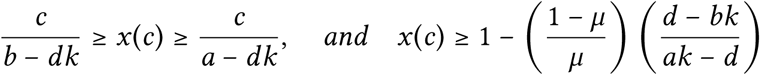

*there exist separating equilibria in which θ*_*H*_ *chooses c*′= 0 *and θ*_*N*_ *chooses c. The level of attention that maximizes the receiver’s payoff satisfies one of these three conditions:*

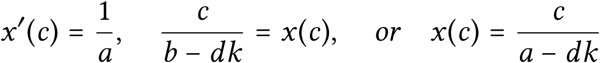

*Proof*. Let type *θ*_*H*_ choose *c*′and type *θ*_*N*_ choose *c*. It is clear that upon observing *c* the receiver will choose *donate* and upon observing *c*′the receiver will choose *decline*. Let the receiver’s best response to *quiet* be *decline*. This holds provided

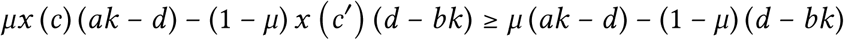

The incentive compatibility constraints for the sender of type *θ* _*H*_ are

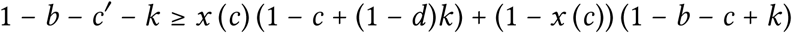

and

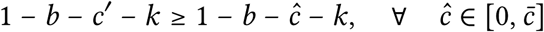

Thus, *c*′must equal 0. Consequently, there is only one non-binding constraint for type *θ*_*H*_, which reduces to

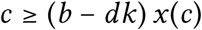

The lone non-binding constraint for type *θ*_*N*_ reduces to

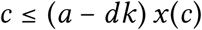

Finally, the receiver’s payoff simplifies to

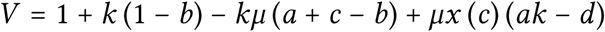

Because of the assumptions about *x* (·), *V* is concave and so the *c* that maximizes *V* is either given by the first order condition with respect to *c* or is a corner solution. Directly,

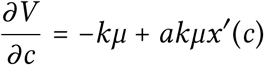

and setting this equal to 0 we obtain

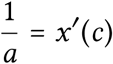

Again, observe that it is the rescaling of the marginal benefit of separating by *x*(*c*) that begets separation. Endogenizing the choice of *x*(·) (as we did for *x*) seems like a worthy enterprise, but one that we will leave for further work.

## 5 Discussion

The primary goal of this paper is to illustrate the counter-intuitive notion that limited attention may facilitate honest communication in situations of conflict. Concisely stated, we discover that inattention rescales the sender’s cost of communication, which lowers the healthy type’s incentive to mimic the needy type. Selective inattention thus can mitigate the effects of an environment unfavorable for communication.

The main results of this paper suggest a number of qualitative and hence testable implications. For instance, as noted in Section 3.1, there are many papers that find non-negligible effects of a distracting environment on an animal’s behavior. Here, we suggest that moderate inattention can encourage honesty. Thus, is there greater honesty among populations in distracting environments? In environments that are extremely distracting, is there a dearth of communication?^18^ Do senders adjust their behavior to account for the receiver’s attention level, i.e., is there more deception when the receiver is likely to be focused?

This paper is related to the line of recent papers in both biology and economics–including Lachmann and Bergstrom (1998); Blume et al. (2007); Rick (2013); Huttegger et al. (2015); Salamanca (2016); and Whitmeyer (2019a)–that highlight that *full transparency* or a perfect communication medium is not generally optimal for the receiver in signaling games. In particular, less than full transparency may beget informative equilibria in situations where there is little to no meaningful communication under full transparency. We establish here that the simple information structures corresponding to inattention are all that is needed for the receiver to achieve her optimum.

Naturally, this paper also fits into the signaling game literature. In addition to the papers cited elsewhere throughout this manuscript, the list of important works includes Yachi (1995), who explores the evolution of honest signaling in a predator-prey interaction; Archetti (2000), who shows that the bright colors of autumn leaves can arise as part of an equilibrium in a signaling game; Huttegger et al. (2014), who explore various dynamical properties of a variety of signaling games; and Sun et al. (2018), who model plant-pollinator interaction as a signaling game. See also the additional discussion of the handicap principle in Számadó (2011) as well as Penn and Számadó (2015), who discuss some of the issues with testing the handicap principle.

Finally, note that the solution concept used throughout this work is a refinement of the standard Nash equilibrium, the subgame perfect equilibrium, and not Evolutionary Stability. However, note that if *x* is fixed and in the interior of *B*, then there is a separating equilibrium that is strict, and thus must therefore be an Evolutionary Stable Strategy in the symmetrized game (see e.g. Cressman (2003)). It might be interesting to explore the dynamic properties of the scenario from this manuscript in greater detail.

## A Omitted Proofs

This appendix contains the proofs and derivations of selected results from the paper.

### A.1 Lemma 3.4 Proof

*Proof*. Suppose the two types of sender separate and let *θ*_*H*_ choose *cry* and *θ*_*N*_ choose *quiet*. Suppose first that 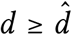, so that *decline* is the receiver’s response to not observing. But then type *θ*_*H*_ ‘s incentive constraint reduces to

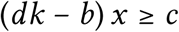

which can never hold. Next, suppose 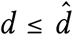, so that *decline* is the receiver’s response to not observing. But then type *θ*_*H*_ ‘s incentive constraint reduces to

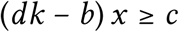

which is always false.

### A.2 Lemma 3.5 Proof

*Proof*. First, it is obvious that there is no equilibrium in which the two types of sender pool on *cry* if *cry* is followed by *decline*.

Next, let 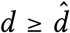 and consider (*quiet, quiet*; *decline*, ·). It is clear that the off-path belief that leaves the equilibrium in greatest jeopardy is that which insists the receiver prefer *donate* upon observing *cry*. Under this “worst” case scenario, the incentive constraint for *θ* _*H*_ is

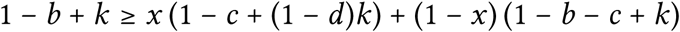

which simplifies to

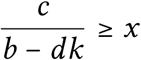

Analogously, the incentive constraint for *θ*_*N*_ reduces to

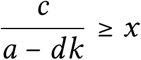

Thus, if the attention parameter *x* ≤ *c*/(*a* − *dk*) then regardless of the receiver’s off-path beliefs, (*quiet, quiet*; *decline*, ·) is an equilibrium. If *x* is above this threshold, then it is clear that an off-path belief that results in the receiver (weakly) preferring *decline* upon observing cry is required.

Now, let 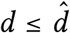. The receiver’s optimal action should she choose not to observe a signal is *donate*. First, we explore whether there is an equilibrium in which the two types of sender pool on *cry*. For *θ* _*H*_ we have

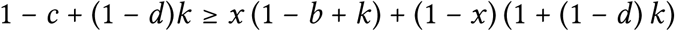

which holds if and only if *x* ≥ *c*/(*b* − *dk*). For *θ*_*N*_ we have

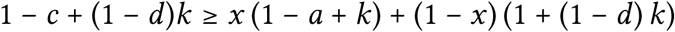

which holds if and only if *x* ≥ *c*/(*a* − *dk*). Note that here we have assigned the receiver’s off-path belief to be such that *decline* is a (weak) best response to *quiet*. This is clearly necessary for the existence of this equilibrium, irrespective of *x*.

Finally, suppose the two types of sender pool on *quiet*. Again, it is clear that this is an equilibrium, regardless of *x* or the off-path beliefs.

### A.3 Proposition 3.8 Proof

This result combines the following two lemmata. Proposition 2.3 is obtained by setting *x* = 1.

#### Lemma A.1.

*Let* 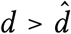.

1. *If* 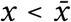, *there exist no equilibria in which θ*_*H*_ *mixes and θ*_*N*_ *chooses cry*.
2. *If* 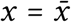, *there exists a continuum of equilibria* (*σ*_*H*_, *cry, decline, donate*), *where*

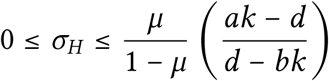
3. *If* 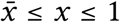, *then there exists an equilibrium* (*σ*_*H*_, *cry, decline, λ*), *where*

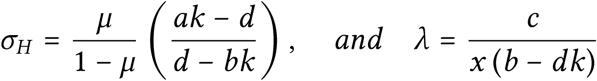

*Proof*. Let *θ*_*H*_ choose mixed strategy *σ*_*H*_ and *θ*_*N*_ choose cry. Naturally, following *quiet* the receiver will strictly prefer *decline*, since she is sure the sender is type *θ*_*H*_. Suppose that the receiver mixes and chooses *donate* with probability *λ* ∈ [0, 1] following *cry*. Then, *θ*_*H*_ ‘s indifference condition requires

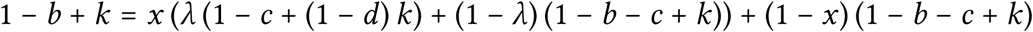

Or,

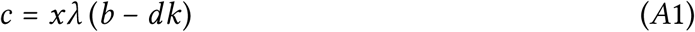

Type *θ*_*N*_ ‘s incentive compatibility condition simplifies to

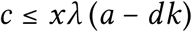

which clearly holds whenever Equation *A*1 does.

Since *c* > 0, *λ* > 0; and so the receiver cannot strictly prefer to choose *decline* following *cry*. First, suppose that *λ* = 1, i.e., that the receiver prefers to choose *donate* after *cry*. Hence,

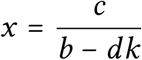

and for the receiver, following an observation of *cry*, the following inequality must hold:

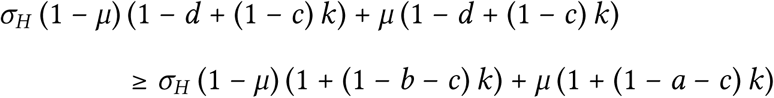

which was derived using Bayes’ law. This reduces to

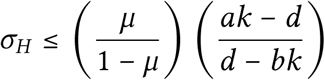

Second, suppose that *λ* ≤ 1 i.e. that the receiver is indifferent between *donate* and *decline* after *cry*. Accordingly,

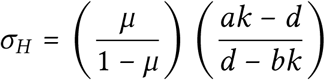

Since *λ* ≤ 1, using *A*1, we must have

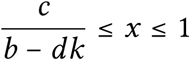

#### Lemma A.2.

*Let* 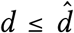. *There exists an equilibrium in which θ*_*H*_ *mixes and θ*_*N*_ *chooses cry if and only if* 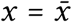.

*Proof*. It is clear that *R* will prefer *decline* following *quiet*, since she knows the sender is type *θ*_*H*_. She prefers *donate* following *cry*, because

1. The prior is such that absent information she weakly prefers to *donate*; and
2. By the martingality of beliefs, if one of her posteriors leaves her more confident that the sender is *θ*_*H*_ (the posterior following *quiet*), then the other posterior must leave her more confident that the sender is *θ*_*N*_.

Type *θ*_*H*_ must be indifferent; hence, which reduces to

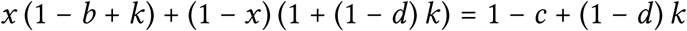

which reduces to

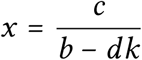

Type *θ*_*N*_ ‘s incentive compatibility condition reduces to

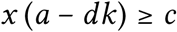

as required.

### A.2 Proposition 3.9 Proof

Just like the proof for Proposition 3.8, this result is the product of two lemmata. Proposition 2.4 is obtained by setting *x* = 1.

#### Lemma A.3.

*Let* 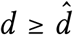. *There exists an equilibrium in which θ*_*N*_ *mixes and θ*_*H*_ *chooses quiet if and only if* 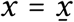.

*Proof*. By the same logic as in Lemma A.2, it is clear that *R* will prefer *decline* following *quiet* and *donate* following *cry. θ*_*N*_ must be indifferent. Hence,

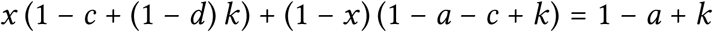

which reduces to

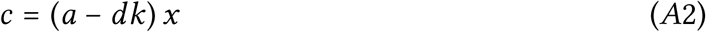

Type *θ*_*H*_ must weakly prefer *quiet*. Thus,

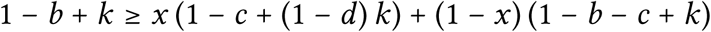

which reduces to

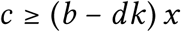

which clearly holds whenever Equation *A*2 does.

#### Lemma A.4.

*Let* 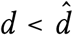.

1. *If x* < 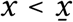, *there exist no equilibria in which θ*_*N*_ *mixes and θ*_*H*_ *chooses quiet*.
2. *If x* = 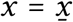, *there exists a continuum of equilibria* (*quiet, σ*_*N*_, *decline, donate*), *where*

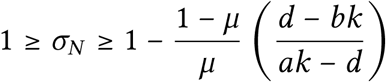
3. *If* 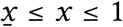, *then there exists an equilibrium* (*quiet, σ*_*N*_, *γ, donate*), *where*

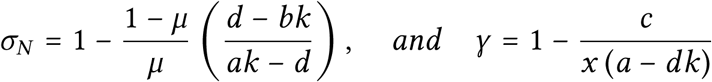

*Proof*. Let *θ*_*N*_ choose mixed strategy *σ*_*N*_ and *θ*_*H*_ choose *quiet*. Naturally, following *cry* the receiver will strictly prefer *donate*. Suppose that the receiver mixes and chooses *donate* with probability *γ*∈ [0, 1] following *quiet*. Then, *θ*_*N*_ ‘s indifference condition requires

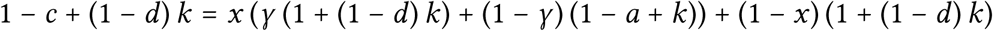

Or,

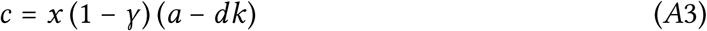

Since *c* > 0,*γ* ≠ 1. Type *θ*_*H*_ ‘s incentive compatibility condition reduces to

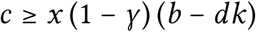

which clearly holds whenever Equation *A*3 does. First, suppose that*γ* = 0, i.e., that the receiver prefers to choose *decline* after *quiet*. Hence,

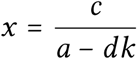

and for the receiver, following an observation of *quiet*, we must have

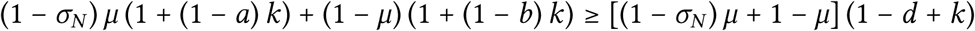

which was derived using Bayes’ law. This reduces to

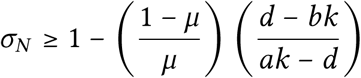

Second, suppose that *γ* ≥ 0, i.e., that the receiver is indifferent between *donate* and *decline* after *quiet*. Accordingly,

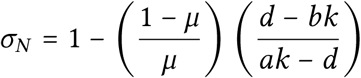

Since *γ* ≥ 0, using Equation *A*3, we must have

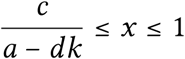

### A.5 Proposition 3.10 Proof

Similarly, this proposition follows from two lemmata. Proposition 2.5 is obtained by setting *x* = 1.

#### Lemma A.5.

*There exist no equilibria in which θ*_*H*_ *mixes and θ*_*N*_ *chooses quiet*.

*Proof*. First, let 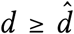. Since *θ*_*H*_ is mixing, he must be indifferent over his pure strategies in support. Hence,

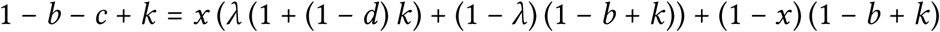

This reduces to

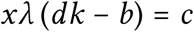

which can never hold.

Second, let 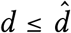. As above, for *θ*_*H*_ we have

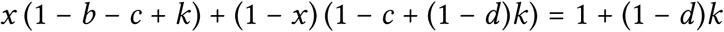

since the receiver prefers *decline* following *cry* and *donate* following *quiet*. This reduces to

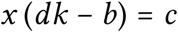

which is always false.

#### Lemma A.6.

*There exist no equilibria in which θ*_*N*_ *mixes and θ*_*H*_ *chooses cry*.

*Proof*. First, let 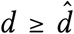. Since *θ*_*N*_ is mixing, he must be indifferent over his pure strategies in support. Hence,

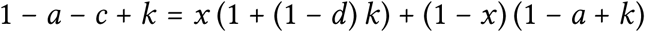

where we have used the fact that following *cry* the receiver will strictly prefer to choose *decline* and following *quiet* the receiver will strictly prefer to choose *donate*. This reduces to *x* (*dk* − *a*) = *c*, which is impossible.

Second, let 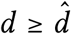. For *θ* _*N*_ we must have

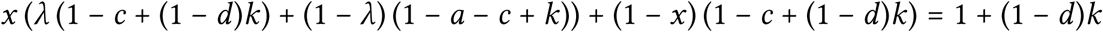

since after *quiet* the receiver strictly prefers to *donate* and after *cry* she chooses *donate* with probability *λ* ∈ [0, 1]. This reduces to *c* = (1 − *λ*) (*dk* − *a*) *x*, which is impossible.

## Footnotes

1 The list of works that have remarked upon this point includes Hurd (1995); Bullock (1997); Bergstrom and Lachmann (1998); Számadó (1999); Lachmann et al. (2001); Bergstrom et al. (2002) and Számadó et al. (2019). See also Penn and Számadó (2020), who provide a comprehensive overview of the handicap principle, its issues, and its relation to empirical findings.

2 Assuming this small but positive search cost is reasonable. As Mock et al. (2011) write, “The central issue…is not whether signals have any detectable cost whatever, which has been demonstrated repeatedly…but ‘whether the costs are sufficient to enforce signal reliability (Searcy and Nowicki (2005)).’ “The monograph they cite (Searcy and Nowicki (2005)) later states in the context of birds signaling through song, “All in all, the bulk of the available evidence suggests that the energy cost of singing, although greater than nothing, is too low for it to put an obvious limit on song output.”

3 Appending an initial stage to a basic signaling game is reminiscent of Számadó (2015), which we discuss in Section 3.1.

4 See, e.g., the “Intuitive Criterion” of Cho and Kreps (1987).

5 The first of these is highlighted by Huttegger and Zollman (2010). Also related is Wagner (2013), who stresses the importance of equilibria of this type in the Spence (1973) signaling game. Kane and Zollman (2015) discover that even when the parameters are such that separation is feasible, evolution is more likely to generate the partially honest signaling characteristic of the mixed-strategy equilibria.

6 To see more discussion of this in communication games see Whitmeyer (2019a) and Whitmeyer (2019b)

7 The modifier “small” is required since we have already imposed several conditions on the values that the parameters may take.

8 Indeed, it is difficult to envision the alternative–absorbing and processing all information from both external and internal stimuli instantaneously.

9 See also Számadó (2018), who introduces competition between senders to the attention-seeking game.

10 We discuss this assumption in Section 3.2.1.

11 This opportunity cost is a theme common to many of papers mentioned in Section 3.1.

12 Other papers that allow for perceptual error, or noise, include Johnstone and Grafen (1992), Johnstone (1994, 1998), Lachmann et al. (2001), and Wiley (2013, 2017). They illustrate that different media may beget different equilibria, and that some media may even foster honest communication impossible in other media.

13 See Whitmeyer (2019a) for a more in depth exposition of this concept.

14 *a* = 1, *b* = 3/8, *c* = 11/64, *d* = 1/8, and *k* = 1/4.

15 I thank an anonymous referee for suggesting this variant of the model.

16 See also the continuous formulation of the Pygmalion game explored in Safley et al. (2020).

17 Alternatively, in the stotting setting, the higher the gazelle jumps, the more likely the predator is to notice how high it jumps but the costlier the jump is in terms of energy expenditure.

18 Recall that when *x* ∈ *A* both sender types are always silent.

## References

Marco Archetti. The origin of autumn colours by coevolution. Journal of Theoretical Biology, 205(4):625 – 630, 2000.

Carl T. Bergstrom and Michael Lachmann. Signalling among relatives. i. is costly signalling too costly? Philosophical Transactions of the Royal Society B: Biological Sciences, 352(1353): 609–617, 1997.

Carl T. Bergstrom and Michael Lachmann. Signaling among relatives. iii. talk is cheap. Proceedings of the National Academy of Sciences, 95(9):5100–5105, 1998.

Carl T. Bergstrom, Szabolcs Számadó, and Michael Lachmann. Separating equilibria in continuous signalling games. Philosophical Transactions of the Royal Society B: Biological Sciences, 357:1595–1606, 2002.

Andreas Blume, Oliver J. Board, and Kohei Kawamura. Noisy talk. Theoretical Economics, 2 (4):395–440, 2007.

Seth Bullock. An exploration of signalling behaviour by both analytic and simulation means for both discrete and continuous models. In Proceedings of the Fourth European Conference on Artificial Life, pages 454–463, Cambridge, Massachusetts, 1997. MIT Press.

Alvin Aaden Yim-Hol Chan, Paulina Giraldo-Perez, Sonja Smith, and Daniel T Blumstein. Anthropogenic noise affects risk assessment and attention: the distracted prey hypothesis. Biology Letters, 6(4):6458 – 6461, 2010.

In-Koo Cho and David M. Kreps. Signaling games and stable equilibria. The Quarterly Journal of Economics, 102(2):179–221, 1987.

Vincent P. Crawford and Joel Sobel. Strategic information transmission. Econometrica, 50(6): 1431–1451, 1982.

Ross Cressman. Evolutionary Dynamics and Extensive Form Games. MIT Press, Cambridge, MA, 2003.

Reuven Dukas. Behavioural and ecological consequences of limited attention. Philosophical Transactions of the Royal Society B: Biological Sciences, 357:1539–1547, 2002.

Reuven Dukas and Alan C. Kamil. The cost of limited attention in blue jays. Behavioral Ecology, 11(5):502–506, 09 2000.

Reuven Dukas and Alan C. Kamil. Limited attention: the constraint underlying search image. Behavioral Ecology, 12(2):192–199, 03 2001.

Drew Fudenberg and David K. Levine. Maintaining a reputation when strategies are imperfectly observed. The Review of Economic Studies, 59(3):561–579, 1992.

Jean-Guy J. Godin and Shelley A. Smith. A fitness cost of foraging in the guppy. Nature, 333: 69–71, 1988.

Peter L. Hurd. Communication in discrete action-response games. Journal of Theoretical Biology, 174(2):217 – 222, 1995.

Simon Huttegger, Brian Skyrms, Pierre Tarrès, and Elliott Wagner. Some dynamics of signaling games. Proceedings of the National Academy of Sciences, 111:10873–10880, 2014.

Simon M. Huttegger and Kevin J. S. Zollman. Dynamic stability and basins of attraction in the sir philip sidney game. Proceedings of the Royal Society B, 277, 2010.

Simon M. Huttegger, Justin P. Bruner, and Kevin J. S. Zollman. The handicap principle is an artifact. Philosophy of Science, 82(5):997–1009, 2015.

Rufus A. Johnstone. Honest signalling, perceptual error and the evolution of ‘all-or-nothing’ displays. Proceedings of the Royal Society B, 256, 1994.

Rufus A. Johnstone. Efficacy and honesty in communication between relatives. The American Naturalist, 152(1):45–58, 1998.

Rufus A. Johnstone and Alan Grafen. Error-prone signalling. Proceedings of the Royal Society B, 248, 1992.

Ulrika Kaby and Johan Lind. What limits predator detection in blue tits (parus caeruleus): posture, task or orientation? Behavioral Ecology and Sociobiology, 54, 2003.

Patrick Kane and Kevin J. S. Zollman. An evolutionary comparison of the handicap principle and hybrid equilibrium theories of signaling. PLOS One, 10(9):1–14, 09 2015.

Jens Krause and Jean-Guy J. Godin. Influence of prey foraging posture on flight behavior and predation risk: predators take advantage of unwary prey. Behavioral Ecology, 7(3): 264–271, 10 1996.

Michael Lachmann and Carl T. Bergstrom. Signalling among relatives: ii. beyond the tower of babel. Theoretical Population Biology, 54(2):146 – 160, 1998.

Michael Lachmann, Szabolcs Számadó, and Carl T. Bergstrom. Cost and conflict in animal signals and human language. Proceedings of the National Academy of Sciences, 98(23):13189–13194, 2001.

David K. Lewis. Convention. A Philosophical Study. Harvard University Press, Harvard, 1969.

Manfred Milinski. A predator’s costs of overcoming the confusion-effect of swarming prey. Animal Behaviour, 32(4):1157 – 1162, 1984.

Douglas W. Mock, Matthew B. Dugas, and Stephanie A. Strickler. Honest begging: expanding from Signal of Need. Behavioral Ecology, 22(5):909–917, 07 2011.

Terry J. Ord. Receiver perception predicts species divergence in long-range communication. Animal Behaviour, 83(1):3 – 10, 2012.

Terry J. Ord and Judy A. Stamps. Alert signals enhance animal communication in “noisy” environments. Proceedings of the National Academy of Sciences, 105(48):18830–18835, 2008.

Kazutaka Ota. Fight, fatigue and flight: narrowing of attention to a threat compensates for decreased anti-predator vigilance. Journal of Experimental Biology, 221(7), 2018.

Dustin J. Penn and Szabolcs Számadó. Why does costly signalling evolve? challenges with testing the handicap hypothesis. Animal Behaviour, 110:e9–e12, 2015.

Dustin J. Penn and Szabolcs Számadó. The handicap principle: how an erroneous hypothesis became a scientific principle. Biological Reviews, 95(1):267–290, 2020.

Douglas G. Richards. Alerting and message components in songs of rufous-sided towhees. Behaviour, 76(3/4):223–249, 1981.

Armin Rick. The benefits of miscommunication. Mimeo, November 2013.

Joshua Safley, Shan Sun, and Jan Rychtár. Dishonest signalling in a variant of pygmalion game. Dynamic Games and Applications, 10:719–731, 2020.

Andrés Salamanca. The value of mediated communication. Mimeo, 2016.

William A. Searcy and Stephen Nowicki. The Evolution of Animal Communication: Reliability and Deception in Signaling Systems. Princeton University Press, 2005.

John Maynard Smith. Honest signalling: the philip sidney game. Animal Behavior, 42:1034–1035, 1991.

Michael Spence. Job Market Signaling. The Quarterly Journal of Economics, 87(3):355–374, 08 1973.

Shan Sun, Michael I. Leshowitz, and Jan Rychtár. The signalling game between plants and pollinators. Scientific Reports, 8(6686), 2018.

Szabolcs Számadó. The validity of the handicap principle in discrete action–response games. Journal of Theoretical Biology, 198(4):593 – 602, 1999.

Szabolcs Számadó. The cost of honesty and the fallacy of the handicap principle. Animal Behaviour, 81(1):3 – 10, 2011.

Szabolcs Számadó. Attention-seeking displays. PLOS ONE, 10(8):1–20, 08 2015.

Szabolcs Számadó. Attention seeking in a spatially explicit game of mate choice and the evolution of dimorphic ornaments. bioRxiv, 2018. URL https://www.biorxiv.org/content/early/2018/01/31/257329.

Szabolcs Számadó, Dániel Czégel, and István Zachar. One problem, too many solutions: How costly is honest signalling of need? PLOS One, 14(1):1–14, 01 2019.

Elliot O. Wagner. The dynamics of costly signaling. Games, 4:163–181, 2013.

Mark Whitmeyer. Bayesian elicitation. ArXiv e-prints, January 2019a. URL https://warxiv.org/abs/1902.00976.

Mark Whitmeyer. In simple communication games, when does ex ante fact-finding benefit the receiver. ArXiv e-prints, January 2019b. URL https://arxiv.org/abs/2001.09387.

R. Haven Wiley. A receiver-signaler equilibrium in the evolution of communication in noise. Behaviour, 150(9/10):9570993, 2013.

R. Haven Wiley. How noise determines the evolution of communication. Animal Behaviour, 124:307 – 313, 2017.

Shigeo Yachi. How can honest signalling evolve? the role of handicap principle. Proceedings: Biological Sciences, 262(1365):283–288, 1995.

Amotz Zahavi. Mate selection—a selection for a handicap. Journal of Theoretical Biology, 53 (1):205 – 214, 1975.

Kevin J. S. Zollman, Carl T. Bergstrom, and Simon M. Huttegger. Between cheap and costly signals: the evolution of partially honest communication. Proceedings of the Royal Society B, 280, 2013.

